# Birth-and-death evolution of the fatty acyl-CoA reductase (FAR) gene family and diversification of cuticular hydrocarbon synthesis in *Drosophila*

**DOI:** 10.1101/509588

**Authors:** Cédric Finet, Kailey Slavik, Jian Pu, Sean B. Carroll, Henry Chung

## Abstract

The birth-and-death evolutionary model proposes that some members of a multigene family are phylogenetically stable and persist as a single copy over time whereas other members are phylogenetically unstable and undergo frequent duplication and loss. Functional studies suggest that stable genes are likely to encode essential functions, while rapidly evolving genes reflect phenotypic differences in traits that diverge rapidly among species. One such class of rapidly diverging traits are insect cuticular hydrocarbons (CHCs), which play dual roles in chemical communications as short-range recognition pheromones as well as protecting the insect from desiccation. Insect CHCs diverge rapidly between related species leading to ecological adaptation and/or reproductive isolation. Because the CHC and essential fatty acid biosynthetic pathways share common genes, we hypothesized that genes involved in the synthesis of CHCs would be evolutionary unstable, while those involved in fatty acid-associated essential functions would be evolutionary stable. To test this hypothesis, we investigated the evolutionary history of the fatty acyl-CoA reductases (FARs) gene family that encodes enzymes in CHC synthesis. We compiled a unique dataset of 200 FAR proteins across 12 *Drosophila* species. We uncovered a broad diversity in FAR content which is generated by gene duplications, subsequent gene losses, and alternative splicing. We also show that FARs expressed in oenocytes and presumably involved in CHC synthesis are more unstable than FARs from other tissues. We suggest that a comparative approach investigating the birth-and-death evolution of gene families can identify candidate genes involved in rapidly diverging traits between species.

## Introduction

Multigene families are important contributors to molecular and organismal evolution. Member genes descend from single founder genes that duplicate, then diverge in sequence (Nei and Rooney 2005). Several models have been proposed to account for how multigene families evolve. For example, the concerted evolution model hypothesizes that all member genes in the family evolve as a unit. This model is capable of explaining aspects of the evolution of clustered ribosomal RNAs (Brown, et al. 1972). In contrast, the birth-and-death model proposes that the members of a gene family evolve independently, meaning that while some members of a gene family are phylogenetically stable, others are unstable and are gained or lost over time by DNA duplications, deletions, and other pseudogenization events (Hughes and Nei 1989; Lynch and Conery 2000; Sjodin, et al. 2007; Plata and Vitkup 2014). Gene repertoire expansion and contraction has been found in diverse gene families such as innate immune genes (Zhang, et al. 2015; Sackton, et al. 2017), plant secondary metabolic genes (Lespinet, et al. 2002; Jiang, et al. 2015; Wang, et al. 2018), developmental transcription factors (Amores, et al. 2004; Tanabe, et al. 2005; Finet, et al. 2012), and snake toxin genes (Dowell, et al. 2016). The accumulated evidence indicates that most gene families evolve according to the birth-and-death model.

It has been suggested that gene birth-and-death could provide insights into the origins of phenotypic novelties (Nei and Rooney 2005; Benton 2015). One example is the cytochrome P450 multigene family (Feyereisen 1999). In animals, all phylogenetically stable P450s encode enzymes that have known endogenous substrates whereas most of the unstable P450s encode enzymes that play roles in xenobiotic detoxification (Thomas 2007; Chung, et al. 2009; Good, et al. 2014). Such observations suggest that members of a gene family with core functions in development and physiology are unlikely to be gained or lost during evolution, whereas members with rapidly-evolving functions between species, such as environmental toxin detoxification, would be gained and lost as species adapt to different habitats (Tatusov, et al. 1997; Rubin, et al. 2000; Hahn, et al. 2007; Thomas 2007; Guo 2013).

Frequent gain and loss of members of a gene family is also apparent in the evolution of the gene families involved in insect chemoreception, such as the gustatory receptor (GR), odorant receptor (OR), and odorant-binding protein (OBP) gene families, which have been shown to expand and contract by birth-and-death evolution (Vieira, et al. 2007; Benton 2015). For instance, in the *D. melanogaster* group, OBP content has evolved more rapidly in the specialist lineages than in their closest generalist relatives (Vieira, et al. 2007). GR and OR repertoires have also been shown to differ considerably between species, with host specialist species losing genes at a much faster rate than their closest generalist sibling species (McBride 2007; McBride, et al. 2007). Together, these data suggest that the genes involved in chemoreception experience rapid evolution among species, and that ecological diversification and natural selection may play major roles in this process.

Among chemoreceptor ligands in insects, short-range or contact pheromones are chemicals that constitute the major signal used in mate recognition between two individuals (Yew and Chung 2015). In many insects, these pheromone components are cuticular hydrocarbons (CHCs) (Howard and Blomquist 2005). Comprised of alkanes, methyl-branched alkanes, and unsaturated hydrocarbons, CHCs form a waxy layer on the cuticle of the insect, where their primary role is probably in maintaining water balance, preventing desiccation due to cuticular water loss (Gibbs 1998). Because of the dual roles that insect CHCs play in both ecological divergence and chemical signaling, these compounds can evolve rapidly among species adapted to living in different environments and habitats (Jallon and David 1987; Chung and Carroll 2015).

The mechanisms underlying the rapid evolution of CHC content are not well understood. The diversity of insect CHCs is shaped by the action of several families of enzymes in specialized cells called oenocytes that are located beneath the insect cuticle (Billeter, et al. 2009). These gene families include fatty acid synthases, desaturases, elongases, and reductases, which make up the ubiquitous fatty acid synthesis pathway in almost all cells. In the oenocytes, a single decarbonylase, *Cyp4g1*, converts some of the products of this pathway into CHCs in *Drosophila* (Qiu, et al. 2012) (fig. 1). Orthologues of *Cyp4g1* has been found in multiple species of insects and have been shown to perform similar functions (Chen, et al. 2016; Yu, et al. 2016; MacLean, et al. 2018). Only a handful of genes encoding enzymes involved in the biosynthesis of CHCs have been identified and characterized so far in *Drosophila* (Chung and Carroll 2015). *mFAS*, which encodes a fatty acyl-CoA synthase expressed in oenocytes, is involved in the production of methyl-branched CHCs (Chung, et al. 2014). The desaturases, desat1 and desat2, play a role in the synthesis of hydrocarbons with at least one double bond at the Z-7 and Z-5 positions, respectively (Dallerac, et al. 2000; Takahashi, et al. 2001), whereas desatF catalyzes the formation of a second double bond in dienes (Chertemps, et al. 2006; Shirangi, et al. 2009). Likewise, eloF is the only elongase known to be involved in the female-specific elongation of long-chain dienes in *D. melanogaster* (Chertemps, et al. 2007). A recent genome-wide association study identified novel fatty acid biosynthesis pathway enzymes that are associated with intraspecific CHC variation in *D. melanogaster*, including three elongases (*CG30008, CG18609* and *CG9458*) and two fatty acyl-coA reductases (FARs) (*CG13091* and *CG10097*) (Dembeck, et al. 2015).

**Fig. 1.**
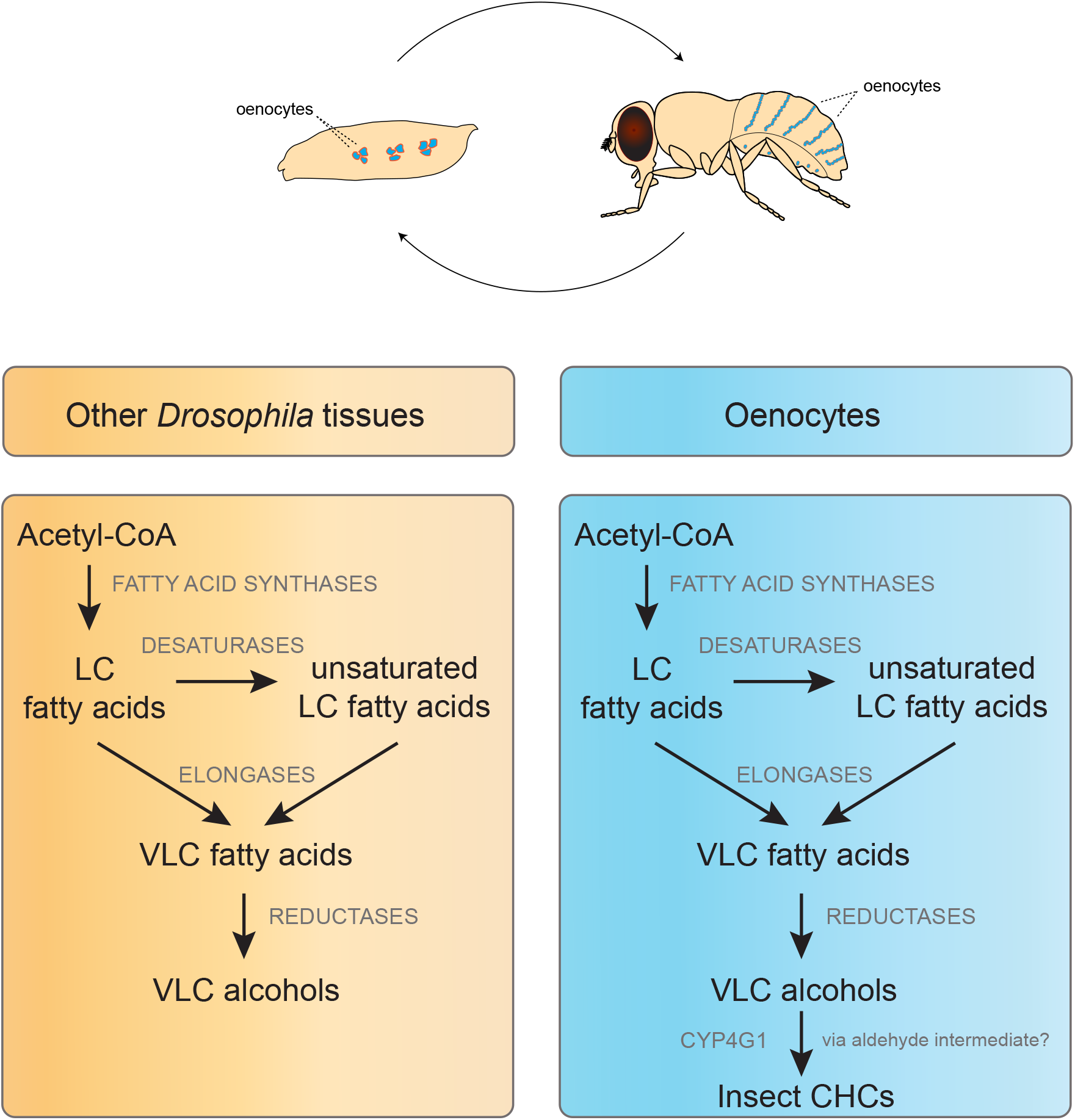
Fatty acid biosynthesis pathway in *Drosophila*. This stepwise process takes place in many cell types, and is catalyzed by several classes of enzymes that generate the diversity of fatty acids found in the organism. Some of these fatty acids are reduced to alcohols by specific reductases. In oenocytes, an additional step consists of the conversion of some of the very long chained (VLC) alcohols to hydrocarbons by a single cytochrome P450, CYP4G1, which exists in all insect genomes sequenced to date.

The majority of the enzymes involved in the synthesis of the CHCs in *Drosophila* are still unknown. The identification of such enzymes has been hampered by experimental difficulties because the CHC pathway has many genes in common with the more pleiotropic fatty acid biosynthesis pathway, and the gene families involved are usually very large (Chung and Carroll 2015). We hypothesized that since CHCs are rapidly evolving traits, the genes underlying their synthesis be rapidly evolving between closely related species. Here, we tested this hypothesis by focusing on the evolution of the FAR gene family that encodes enzymes catalyzing the reduction of acyl-CoA to alcohols and aldehydes (Riendeau and Meighen 1985; Cinnamon, et al. 2016) (fig. 1). We show that the FAR gene family evolves following a birth-and-death model. We took advantage of differential molecular evolutionary features between stable and unstable FAR lineages to identify the FARs that are likely to be involved in core functions (fatty acid biosynthesis) and those likely to be involved in rapidly-evolving functions between species (CHC biosynthesis).

## Results

### Combined phylogenetic and microsynteny analyses identify stable and unstable members of the FAR gene family in *Drosophila*

To determine which members of the FAR gene family are evolutionarily stable or unstable, we employed a phylogenomic approach based on an unprecedented sampling of FAR sequences. We applied an exhaustive BLAST similarity search to the 12 available full *Drosophila* genomes. We found that the number of FARs in each of the 12 sequenced *Drosophila* genomes ranged from 14 to 21 (table 1). We then performed maximum-likelihood and Bayesian phylogenetic reconstruction on our 200-FAR dataset (supplementary fig. S1). The resulting tree clarifies the number of main FAR lineages within the *Drosophila* genus. The FAR sequences split into 18 main clades (fig. 2A). Out of these 18 clades, three clades originated through gene duplications specific to the *melanogaster* group such as the duplication leading to the clades *CG13091* and *CG10097*, and the one leading to the clades *CG17562, CG14893*, and *CG17560* (fig. 2B). After removal of these lineage-specific FAR clades, it is reasonable to infer that the last common ancestor of the extant *Drosophila* genus possessed at least 15 FAR genes.

**Table 1.** FAR gene content in the 12 sequenced *Drosophila* genomes. The number of FARs ranges from 14 to 21, with 17 genes in the model species *D. melanogaster*.

| Species | FAR Content | Genome | Previous estimates |
| --- | --- | --- | --- |
| *Drosophila melanogaster* | 17 | v. 6.06 | 13 (Lassance et al. 2010); 15 (Eirín-López et al. 2012) |
| *Drosophila simulans* | 17 | v. 2.01 | This study |
| *Drosophila sechellia* | 17 | v. 1.3 | This study |
| *Drosophila yakuba* | 17 | v. 1.04 | This study |
| *Drosophila erecta* | 17 | v. 1.04 | This study |
| *Drosophila ananassae* | 17 | v. 1.04 | This study |
| *Drosophila pseudoobscura* | 21 | v. 3.03 | This study |
| *Drosophila persimilis* | 20 | v. 1.3 | This study |
| *Drosophila willistoni* | 15 | v. 1.04 | This study |
| *Drosophila mojavensis* | 14 | v. 1.04 | This study |
| *Drosophila virilis* | 20 | v. 1.03 | This study |
| *Drosophila grimshawi* | 14 | v. 1.3 | This study |

**Fig. 2.**
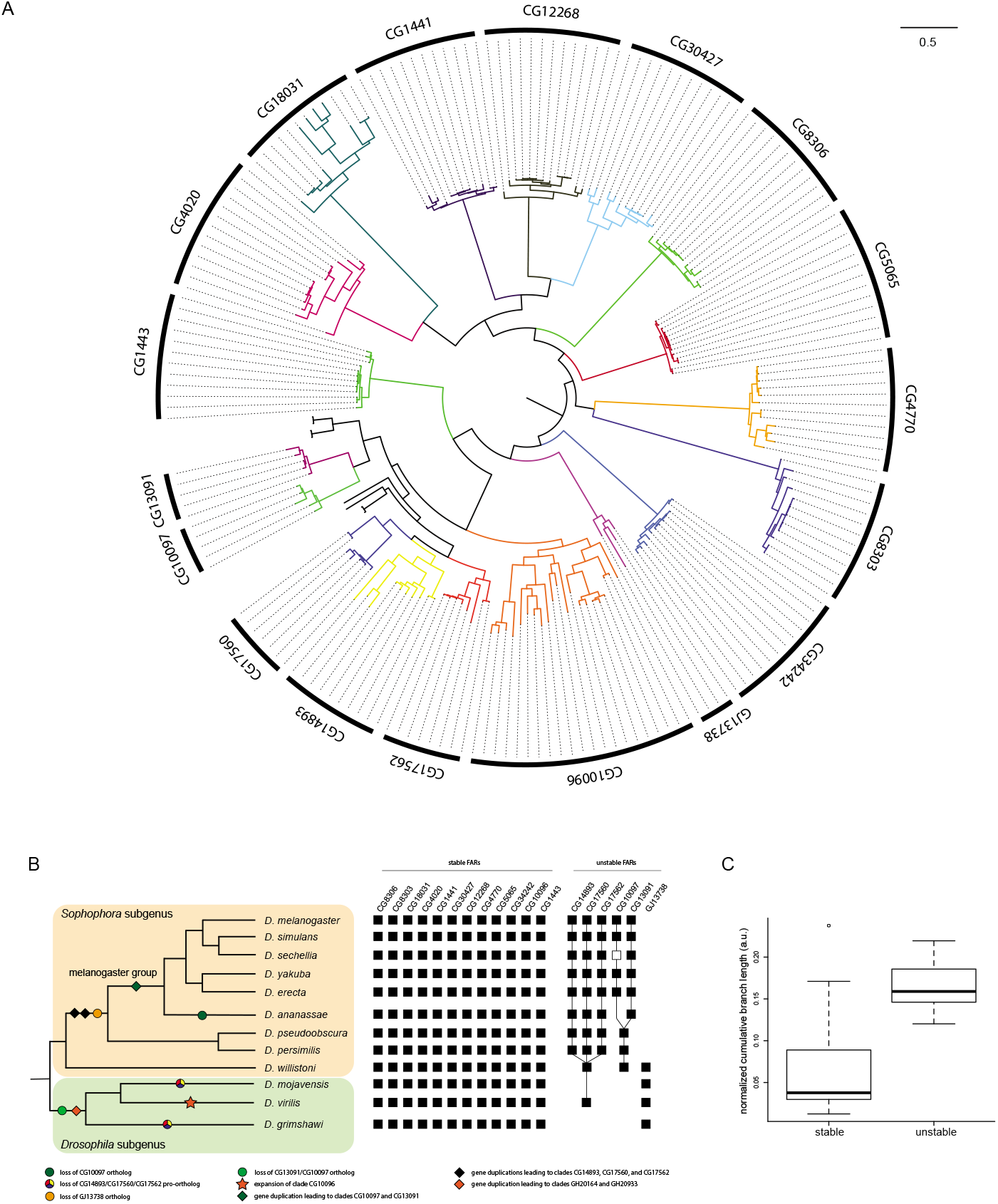
Phylogeny and evolution of FAR genes in the *Drosophila* genus. (A) Phylogram of the 200-taxon analyses. RAxML maximum-likelihood analyses and PhyloBayes Bayesian analyses were conducted under the LG and the GTR+Γ model, respectively. Support values are shown in supplementary figure X). Scale bar indicates number of changes per site. (B) Molecular events underlying the origin and diversification of the FAR repertoire. Filled squares indicate the presence of a gene, and open squares indicate pseudogenes. (C) Comparison of cumulative branch lengths between stable and unstable FAR clades.

Our inference of the number of clades is also well supported by the study of microsynteny conservation between species. Each clade includes orthologous sequences whose relative genomic location is conserved between the 12 *Drosophila* genomes (supplementary fig. S2). We identified a set of 12 stable FARs with at least one copy in all 12 genomes (fig. 2). Conversely, we identified a set of unstable FAR clades that show variable gene content among *Drosophila* species. Moreover, unstable FAR and stable FAR genes have very distinct branch lengths, with longer branch lengths for unstable FARs (t-test: df=13, P=0.003; fig. 2C), suggesting that unstable FAR genes evolve faster than stable FAR genes.

### The Drosophila FAR repertoire evolved through multiple gene duplication events and independent gene losses

Several independent gene losses have occurred during the evolution of the FAR gene family. For example, FAR genes of the clade *CG13091*/*CG10097* are absent from the entire subgenus *Drosophila* (grimshawi, mojavensis, and virilis groups), and the *CG10097* orthologue is absent from *D. ananassae* (fig. 2B). Moreover, the *CG10097* orthologue in *D. sechellia (DsecGM26015)* shows clear features of pseudogenization such as a fast evolution rate (suppl. Fig.1, table 2) and a 16bp-deletion that results in a truncated putative transcript. In contrast, genes of the clade *CG13091*/*CG10097* have been duplicated and retained in most species of the subgenus *Sophophora* (except for the ananassae group).

**Table 2.** Statistics for the branch-site test of positive selection. These values were obtained by applying the branch-site test implemented in the package codeml of PAML. df: degree of freedom, lnL: log likelihood score, −2ΔlnL: likelihood ratio test.

| foreground branch | null model | | alternative model | | $-2\Delta\ln L$ | df | p-value |
| --- | --- | --- | --- | --- | --- | --- | --- |
| | parameters | $\ln L$ | parameters | $\ln L$ | | | |
| *DanaGF17060* | $p_0 = 0.561$<br>$p_1 = 0.126$<br>$p_{2a} = 0.255$<br>$p_{2b} = 0.058$<br>$\omega_0 = 0.124$<br>$\omega_1 = 1$<br>$\omega_2 = 1$ | -11225.209 | $p_0 = 0.744$<br>$p_1 = 0.167$<br>$p_{2a} = 0.073$<br>$p_{2b} = 0.016$<br>$\omega_0 = 0.124$<br>$\omega_1 = 1$<br>$\omega_2 = 11.202$ | -11219.170 | 12.078 | 1 | $5.1 \times 10^{-4}$ |
| *DanaGF17063* | $p_0 = 0.709$<br>$p_1 = 0.167$<br>$p_{2a} = 0.1$<br>$p_{2b} = 0.024$<br>$\omega_0 = 0.123$<br>$\omega_1 = 1$<br>$\omega_2 = 1$ | -11226.499 | $p_0 = 0.762$<br>$p_1 = 0.174$<br>$p_{2a} = 0.051$<br>$p_{2b} = 0.012$<br>$\omega_0 = 0.126$<br>$\omega_1 = 1$<br>$\omega_2 = 14.503$ | -11223.058 | 6.884 | 1 | $8.7 \times 10^{-3}$ |
| *DsecGM26015* | $p_0 = 0.378$<br>$p_1 = 0.063$<br>$p_{2a} = 0.479$<br>$p_{2b} = 0.080$<br>$\omega_0 = 0.116$<br>$\omega_1 = 1$<br>$\omega_2 = 1$ | -4554.441 | $p_0 = 0.800$<br>$p_1 = 0.146$<br>$p_{2a} = 0.045$<br>$p_{2b} = 0.009$<br>$\omega_0 = 0.117$<br>$\omega_1 = 1$<br>$\omega_2 = 257.742$ | -4529.358 | 50.167 | 1 | $1.4 \times 10^{-12}$ |
| *DperGL27182/DpseGA32357* | $p_0 = 0.766$<br>$p_1 = 0.136$<br>$p_{2a} = 0.083$<br>$p_{2b} = 0.015$<br>$\omega_0 = 0.125$<br>$\omega_1 = 1$<br>$\omega_2 = 1$ | -4569.562 | $p_0 = 0.815$<br>$p_1 = 0.147$<br>$p_{2a} = 0.032$<br>$p_{2b} = 0.006$<br>$\omega_0 = 0.125$<br>$\omega_1 = 1$<br>$\omega_2 = 13.669$ | -4565.092 | 8.941 | 1 | $2.8 \times 10^{-3}$ |
| *DvirGJ21443* | $p_0 = 0.587$<br>$p_1 = 0.222$<br>$p_{2a} = 0.139$<br>$p_{2b} = 0.052$<br>$\omega_0 = 0.143$<br>$\omega_1 = 1$<br>$\omega_2 = 1$ | -14488.933 | $p_0 = 0.682$<br>$p_1 = 0.256$<br>$p_{2a} = 0.045$<br>$p_{2b} = 0.017$<br>$\omega_0 = 0.144$<br>$\omega_1 = 1$<br>$\omega_2 = 15.825$ | -14480.535 | 16.795 | 1 | $4.2 \times 10^{-5}$ |
| *DvirGJ22672/22673/26512* | $p_0 = 0.679$<br>$p_1 = 0.260$<br>$p_{2a} = 0.044$<br>$p_{2b} = 0.017$<br>$\omega_0 = 0.144$<br>$\omega_1 = 1$<br>$\omega_2 = 1$ | -14491.793 | $p_0 = 0.703$<br>$p_1 = 0.267$<br>$p_{2a} = 0.022$<br>$p_{2b} = 0.008$<br>$\omega_0 = 0.144$<br>$\omega_1 = 1$<br>$\omega_2 = 8.945$ | -14488.424 | 6.739 | 1 | $9.4 \times 10^{-3}$ |

Another case of FAR gene loss is in the clade *GJ13738*, which is restricted to the entire subgenus *Drosophila* and the *willistoni* group. When mapped onto the *Drosophila* species tree, the distribution of the clade *GJ13738* suggests a unique loss event in the subgenus Sophophora after the divergence of the *willistoni* group (fig. 2B). A third example of FAR gene loss is in the ancestral clade *CG14893/CG17560/CG17562* which is restricted to the subgenus *Sophophora* and *D. virilis*. We find two independent losses of this clade in the lineage leading to *D. mojavensis* and *D. grimshawi*, respectively (fig. 2B). Conversely, this clade went through three successive rounds of gene duplications within the *Sophohora* subgenus.

### Signatures of positive selection associated with repeated duplication events

The most striking example of FAR content expansion is of the *CG10096* orthologues in *D. virilis* (fig. 2A and 2B). Eight copies have been identified in the genome of *D. virilis*. This specific expansion contributes to a higher FAR content in *D. virilis* (table 1), as well as an increase of the cumulative branch length for the *CG10096* clade (see the outlier dot, fig. 2C). This latter observation could result from a faster rate of molecular evolution due to relaxed selective pressure. We tested this hypothesis by searching for any signatures of positive selection in the *CG10096* clade. We did detect several branches and sequences of *D. virilis* under positive selection (table 2). We also noted expansion of the *CG14893* clade in *D. ananassae* to three copies. We detect signatures of positive selection in two of these paralogues (table 2).

### Evolution of putative new substrate specificity: the unique case of clade *CG30427*

We have shown that gene duplication has played a major role in the diversification of the FAR family. We also observed an interesting case of alternative splicing affecting FAR diversity in the CG30427 clade. In *D. melanogaster*, the gene *CG30427* gene produces three main classes of transcripts (fig. 3). Surprisingly, the *CG30427* transcript variants have a highly conserved exon/intron structure, and encode similar protein isoforms. These observations suggest that the gene *CG30427* could have evolved by serial duplication of exons 3-6 leading to repetition of the structural domains. Independent evolution (e.g., mutation) of the repeated exons, as well as the establishment of alternative splicing, could have subsequently generated distinct, but still comparable, isoforms. Notably, one of the predicted substrate-binding sites shows an amino acid difference between isoforms A and C (**M**ethionine) and isoform B (**V**aline). This may be significant because in *Arabidopsis thaliana*, the two enzymes FAR5 and FAR8 are 85% identical at the amino acid level, but they possess distinct substrate specificities for 18:0 or 16:0 acyl chain lengths, respectively (Chacón et al, 2013). Moreover, it has been recently shown that just two individual amino acid substitutions (L355A and M377V) explain most of the difference in substrate specificity between FAR5 and FAR8 (Chacón et al, 2013). Although there is no direct biochemical evidence available yet, we infer that the *D. melanogaster CG30427* transcript variants probably encode functionally distinct isoforms.

**Fig. 3.**
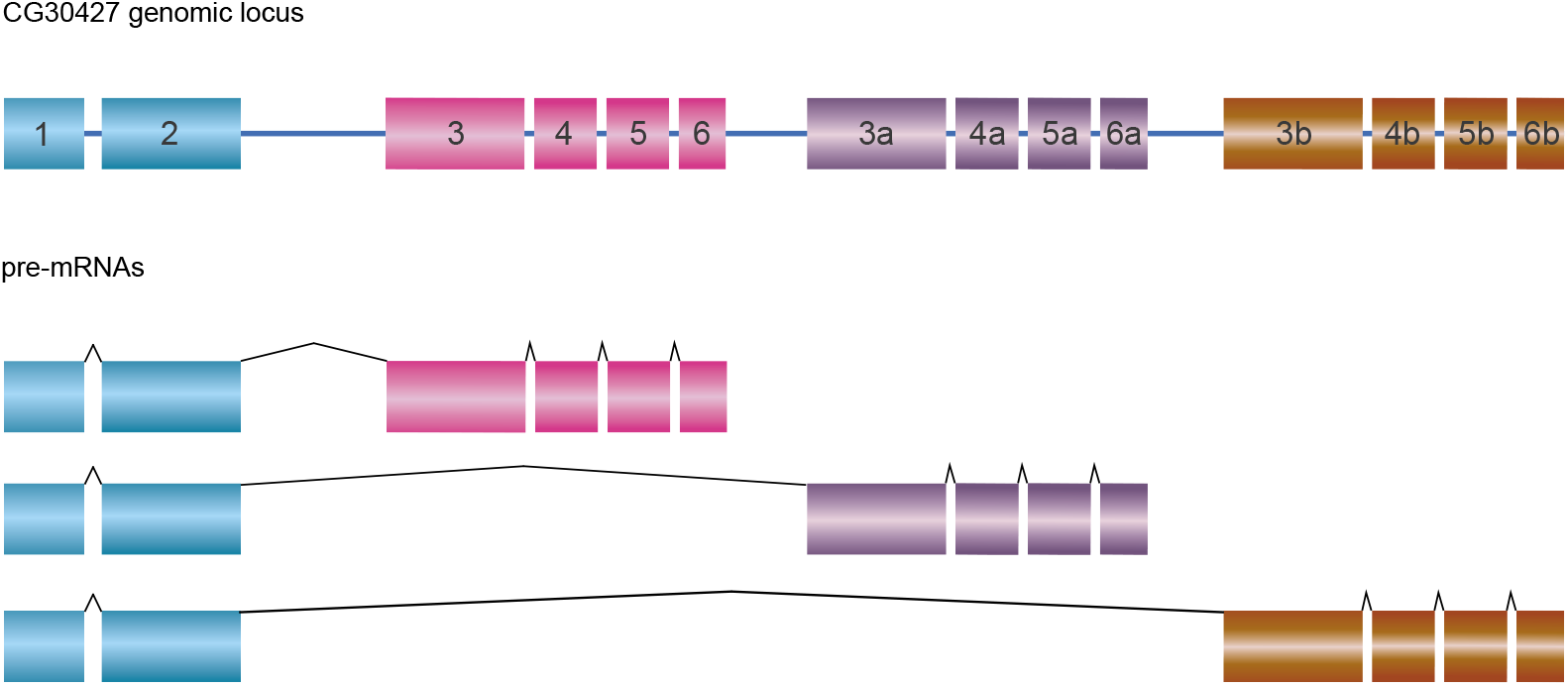
Genomic structure and isoforms of the gene *CG30427* in *D. melanogaster*. Exons 3-6 underwent duplication in tandem and do exist in three different copies. The combination of the three different “cassettes” with the single exons 1-2 leads to three different isoforms.

### Unstable FARs are mostly expressed in the oenocytes, the site of CHC biosynthesis

The biosynthesis of fatty acyl-CoA takes place in many tissues in the fly (Jaspers, et al. 2014), but the biosynthesis of CHCs specifically occurs in the oenocytes (Billeter, et al. 2009; Wicker-Thomas, et al. 2015) (fig. 1). To determine which FARs may be involved in CHC production, we identified FARs expressed in the oenocytes by *in situ* hybridization in *D. melanogaster*. Using DIG-labeled RNA probes for 16 FARs (all except *CG4770*), we performed *in situ* hybridization on both mixed stage embryos and dissected adult abdomens. We detected expression of four FARs in oenocytes from adults. Three of these are evolutionarily unstable: *CG13091* (male expressed), *CG10097* (male expressed), and *CG17560* (expressed in both sexes). The only evolutionarily stable FAR expressed in oenocytes is *CG4020* (female expressed) (fig. 4). *In situ* hybridization in embryos showed only *CG17562* and *CG18031 (FarO)* are expressed in embryonic oenocytes (fig. 4), while the other FARs are expressed in other tissues such as the salivary glands and the tracheal system (supplementary fig. S3). No FAR is expressed in both embryonic and adult oenocytes.

**Fig. 4.**
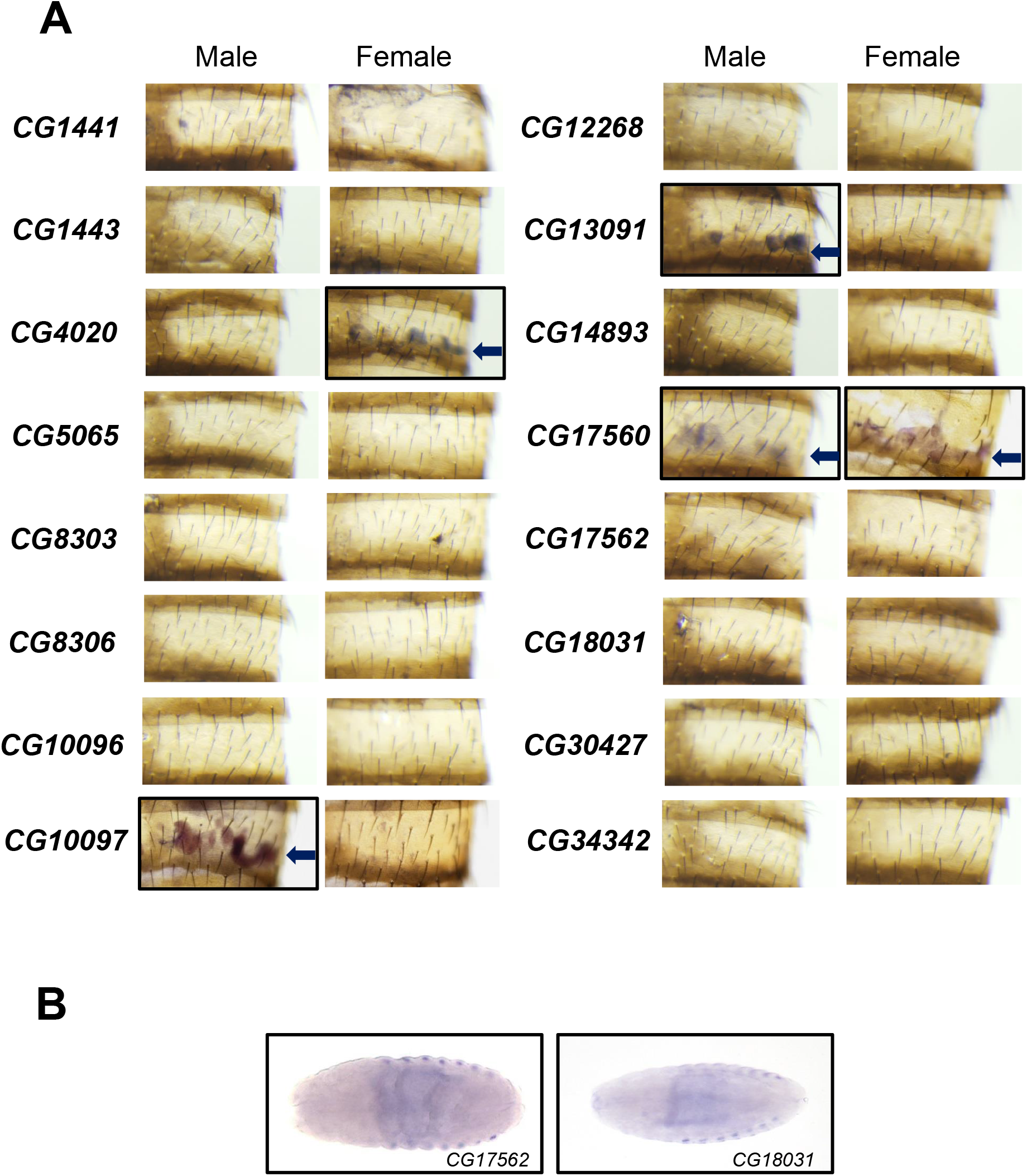
Expression of FARs in *D. melanogaster* adult cuticle and embryos. (A) *In situ* hybridizations of FARs in abdominal cuticle show that only four FARs are expressed in adult oenocytes (see arrows). (B) *In situ* hybridizations of the two FARs *CG17562* and *CG18031* that are expressed in embryonic oenocytes.

### Stable FARs are likely to have essential functions

To determine if the loss of a FAR impacts viability, we used the ubiquitous tubulin-GAL4 driver and UAS-RNA interference (RNAi) to knock down each FAR individually. We found that RNAi knockdown of nine out of the 12 stable FARs (75%) led to mortality while only one out of five unstable FARs (20%) was essential for viability (table 3). These results demonstrate that the majority of stable FARs are essential for viability and confirm previous work on two specific FARs including the *CG1443* gene (wat), which is expressed in the trachea and involved in gas filling of the tracheal tubes during *Drosophila* embryogenesis (Jaspers, et al. 2014), and the *CG18031* gene (*FarO*), which plays a key role in preventing excessive oenocyte cell growth (Cinnamon, et al. 2016). In contrast, we deduce that most of the unstable FARs are involved in nonessential functions that evolve rapidly between species.

**Table 3.** RNAi knockdown of individual FAR genes using the ubiquitous tubulin-GAL4 driver. Most of the evolutionarily stable members of this gene family are essential for development (lethal when knocked down by RNAi), while most of the evolutionarily unstable members of this gene family are involved in non-developmental processes (flies still viable when these genes are knocked down by RNAi).

| Phylogenetic stability | Gene name | RNAi phenotype |
| --- | --- | --- |
| stable | CG1443 | lethal |
|  | CG4020 | lethal |
|  | CG4770 | lethal |
|  | CG5065 | lethal |
|  | CG8303 | lethal |
|  | CG8306 | lethal |
|  | CG10096 | lethal |
|  | CG12268 | lethal |
|  | CG34342 | lethal |
|  | CG1441 | viable |
|  | CG18031 | viable |
|  | CG30427 | viable |
| unstable | CG10097 | viable |
|  | CG13091 | viable |
|  | CG14893 | viable |
|  | CG17562 | viable |
|  | CG17560 | lethal |

## Discussion

Using a combination of bioinformatics and reverse genetics, we have conducted a comprehensive study of the FAR gene family in the genus *Drosophila*. We have shown that five out of 17 FARs found in the *D. melanogaster* genome are evolutionary unstable. Most of these unstable FARs are expressed in the oenocytes, the site of CHC biosynthesis in *D. melanogaster*, compared to only one of the 12 stable FARs. Our functional RNAi experiments demonstrate that most stable FARs carry out have functions crucial for viability, whereas silencing most unstable FARs do not lead to lethality. These data suggest that the gain and loss of unstable FARs can alter CHC diversity without affecting insect viability. Comparison of cumulative branch lengths between stable and unstable FAR clades showed that unstable FARs undergo more rapid sequence evolution compared to stable FARs. Taken together, we suggest that FAR genes involved in CHC synthesis are likely to be evolutionary unstable and evolve faster than other FARs. These results appear to support an important but largely untested tenet of the birth-and-death model of gene families (Nei and Rooney 2005; Eirin-Lopez, et al. 2012) -- that stable members of gene families often encode genes with core functions involved in viability while unstable members often encode genes involved in non-viable and rapidly evolving functions (Thomas 2007).

However, the birth and death of fatty acyl-CoA biosynthesis gene family members is not the only mechanism underlying the rapid divergence of CHCs between species. The fatty acyl-coA desaturase (desat) gene family is another gene family involved in the synthesis of CHCs. Three desats, *(desat1, desat2* and *desatF)* have been experimentally shown to be involved in CHC synthesis in *Drosophila* (Dallerac, et al. 2000; Takahashi, et al. 2001; Chertemps, et al. 2006). *desat1* is an evolutionary stable gene which has pleiotropic functions in *D. melanogaster* (Bousquet, et al. 2012), while *desat2* was lost in *D. erecta*, and the *desatF* lineage went through several rounds of gene duplication and subsequent specific gene losses (Fang, et al. 2009; Keays, et al. 2011). However, regulatory changes that affect oenocyte expression, as well as transition from monomorphic to dimorphic oenocyte expression (and its reversion) of *desatF*, account for CHC divergence caused by this gene as well (Shirangi, et al. 2009). *Cis*-regulatory changes in other fatty acyl-CoA biosynthesis genes has also been shown to be involved in CHC divergence between *Drosophila* species. These include *cis*-regulatory changes in *mFAS* (a fatty acid synthase) expression between two closely related Australian *Drosophila* species (Chung, et al. 2014), as well as a recent discovery that tissue-specific *cis*-regulatory changes affect the expression of *eloF*, a fatty acid elongase, leading to CHC divergence and mating inhibition in *D. simulans* and *D. sechellia* (Combs, et al. 2018). Based on the evidence obtained to date, we suggest that the birth-and-death of fatty acyl-CoA biosynthesis genes, as well as *cis*-regulatory evolution, accounts for the majority of CHC divergence in *Drosophila*. We note that no example of coding changes has been shown thus far to account for CHC divergence.

Differences in gene family content allow the diversification and ecological adaptation of many different species (Demuth and Hahn 2009; Żmieńko, et al. 2014; Carretero-Paulet, et al. 2015). The advancement of sequencing technologies has led to sequencing and availability of more than 100 insect genomes (Yin, et al. 2015) with a few thousand more being proposed (Consortium 2013). This includes closely related species such as 16 *Anopheles* mosquito genomes (Neafsey, et al. 2015). The birth-and-death evolution model could be used to identify genes involved in rapidly evolving traits between species such as CHC synthesis. Because CHCs are involved in premating isolation between many closely related insect species, identification of evolutionarily unstable genes may also shed light on the speciation and radiation of such groups.

## Materials and Methods

### Fly stocks

The *Canton-S* strain or the *Xout* strain was used as the wild-type *D. melanogaster* strain for *in situ* hybridization. All RNAi lines were obtained from the VDRC (Vienna *Drosophila* RNAi center)(Dietzl, et al. 2007). The tubulin-GAL4 (Bloomington Stock #5138) strain was obtained from the Bloomington *Drosophila* Stock Center. All flies were maintained at room temperature on standard Bloomington recipe *Drosophila* food.

### Data collection

Fatty acyl-CoA reductase genes were identified in 12 complete *Drosophila* genomes by tBLASTn using a set of selected *D. melanogaster* sequences as a probe. *Drosophila ananassae, D. erecta, D. grimshawi, D. melanogaster, D. mojavensis, D. persimilis, D. pseudoobscura, D. sechellia, D. simulans, D. virilis, D. willistoni*, and *D. yakuba* genomes were retrieved from the FlyBase website (http://flybase.org). We experimentally checked the sequence of some FARs by PCR, especially when we started to use early versions of the released genomes. We completed the sequence of the *D. yakuba* transcript *FarO (DyakGE28152)*, and deposited the new sequence in the EMBL database (accession number LT996250). The sequence alignments and tree files are downloadable from Dryad (XXX).

### Phylogenetic analyses

Amino acid sequences were aligned with MUSCLE (Edgar 2004), manually adjusted, and selected blocks were used for phylogenetic reconstruction. Maximum-likelihood searches were performed using RAxML 7.3.5 (Stamatakis 2006), which allows efficient maximum-likelihood analyses of large data sets. All searches were completed under the LG substitution matrix with final likelihood evaluation using a gamma distribution. 100 bootstrap replicates were conducted for support estimation. In addition, we used the PhyloBayes 3.3 program (Lartillot, et al. 2009), which implements a GTR+Γ model. We ran two independent chains for at least 21,817 cycles, and discarded the first 5,000 cycles as burn-in. Run parameters of the chains were also plotted together against time to check appropriate convergence and chain mixing.

### Detection of positive selection

Candidate genes were tested for signatures of positive selection based on the ratio ω = d_N_/d_S_ (nonsynonymous/synonymous substitution rates) using the program codeml of PAML v.4.8 (Yang 2007). We compared the null model (ω fixed to 1) to the alternative branch-site model that allows some sites to have an ω greater than one in some branches. The two models were compared using a likelihood ratio test (LRT). A p-value < 0.05 means that the model with positive selection better explains the data. The codon alignment, used as input in PAML, was generated using the software PAL2NAL (Suyama, et al. 2006).

### Statistical analysis

The cumulative branch length (CBL) per clade was calculated by adding all branch lengths within a clade. Branch lengths were obtained as outputs of RAxML software (Stamatakis 2006). To take into account differences in number of sequences per clade, we calculated the normalized CBL, that is, the value of CBL/number of FAR sequences per clade. We compared the CBL means between stable and unstable FARs using a t-test as the CBL followed a normal distribution. Statistical tests and graphics were performed using R statistics package version 3.5.0 (the R Project for Statistical Computing, www.r-project.org, last accessed 2018 April 27).

### In situ hybridization in embryos and adult oenocytes

*In situ* hybridization of oenocytes of embryos or four- to five-day old adults was performed with RNA probes as described previously (Shirangi, et al. 2009). Probes were made from mixed-sex five-day old adult cDNA using the primers listed in Table S1.

### RNAi experiments

To determine if a given reductase was important for the viability of the fly, UAS-RNAi strains were individually crossed to tubulin-GAL4/TM3 *Sb*, resulting in RNAi knockdown in a ubiquitous pattern as previously described (Chung, et al. 2009). Reciprocal crosses were performed at 25°C. The sex and phenotype of emerging adults were scored. Stubble bristles were used to indicate the presence of the TM3, *Sb* chromosome in progeny, and therefore the absence of the tubulin-GAL4 chromosome. A specific reductase was scored as having an essential function if only flies carrying the TM3, *Sb* chromosome emerged from the cross (*i.e*., ubiquitous RNAi of the FAR resulted in lethality).

## Author contribution

C.F. performed the phylogenetic and genomic analyses. K.S, J.P and H.C. performed fly crosses and *in situ* hybridization. C.F., J.G.M, S.B.C. and H.C. conceived the project. C.F. and H.C. wrote the paper. All authors contributed to the drafting of the manuscript.

## Acknowledgements

S.B.C. is a Howard Hughes Medical Institute investigator. H.C. is supported by MSU AgBioresearch umbrella project MICL02522. Stocks obtained from the Bloomington Drosophila Stock Center (NIH P40OD018537) were used in this study. We acknowledge Jocelyn Millar (University of California, Riverside) for discussion during the course of this project and on the manuscript.

## Supplementary data

**Fig. S1.**
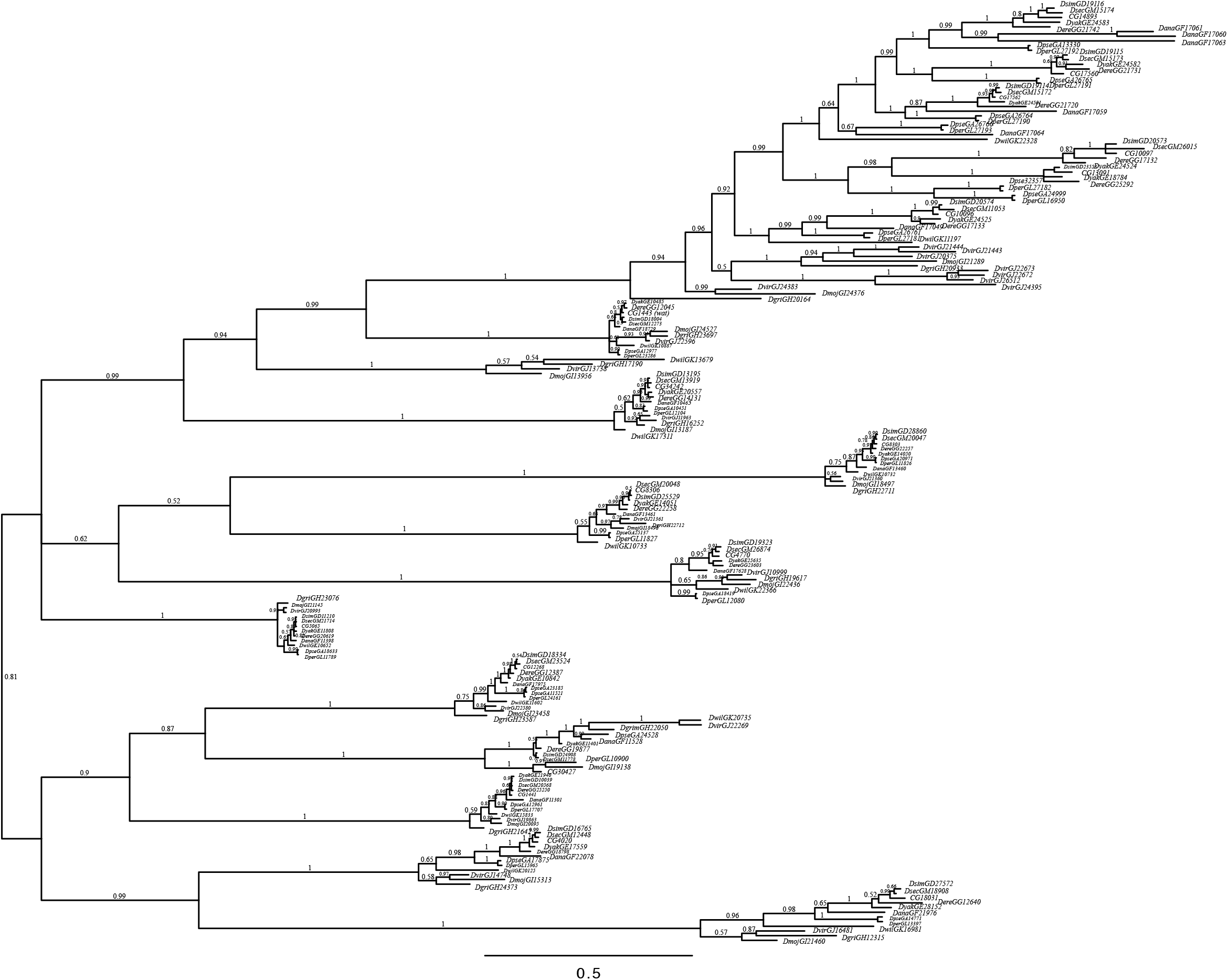
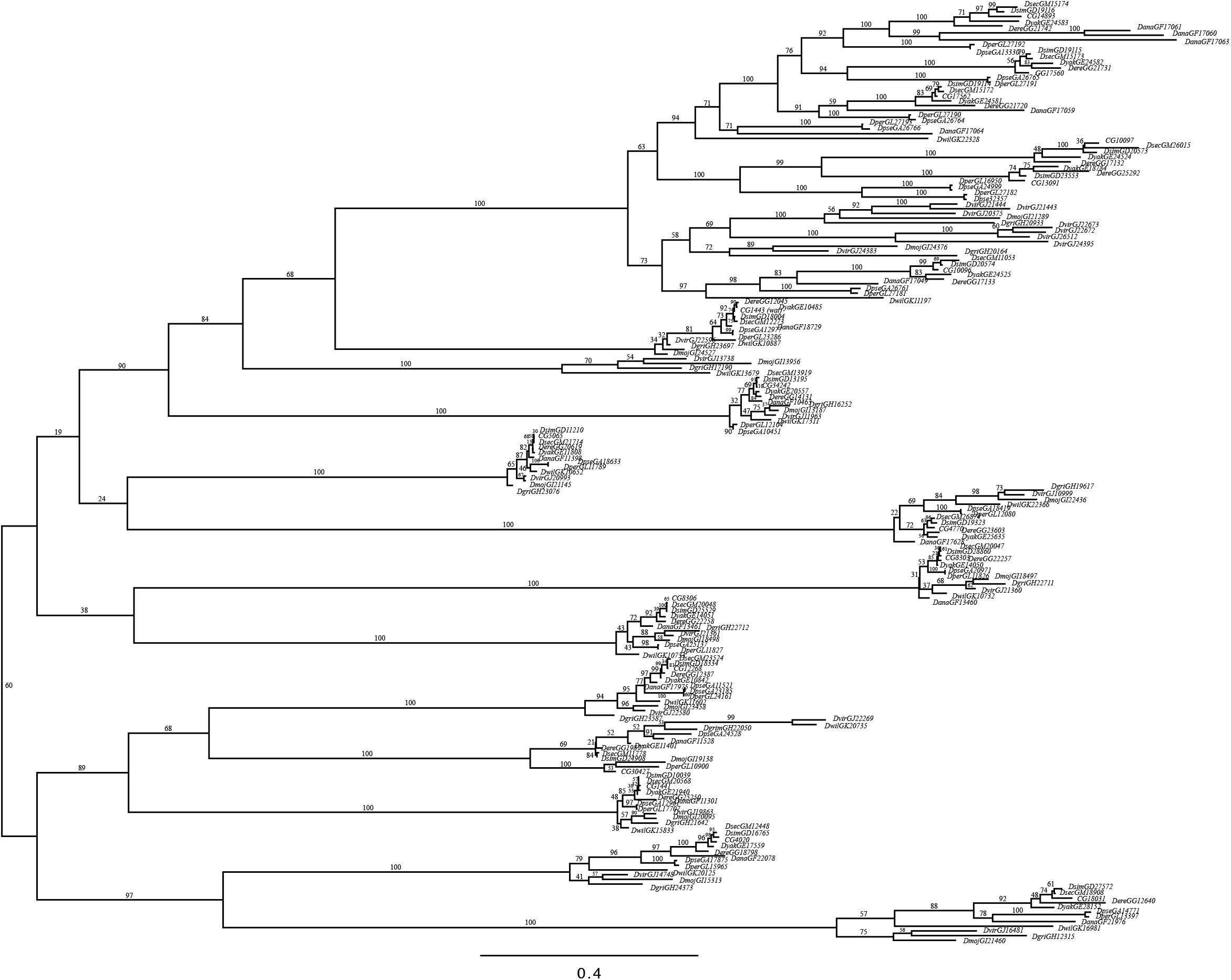
Phylograms of the 200-taxon analyses. RAxML maximum-likelihood analyses and PhyloBayes Bayesian analyses were conducted under the LG and the GTR+Γ model, respectively. Support values obtained after 100 bootstrap replicates and Bayesian posterior probabilities are show for all branches. Scale bar indicates number of changes per site.

**Fig. S2.**
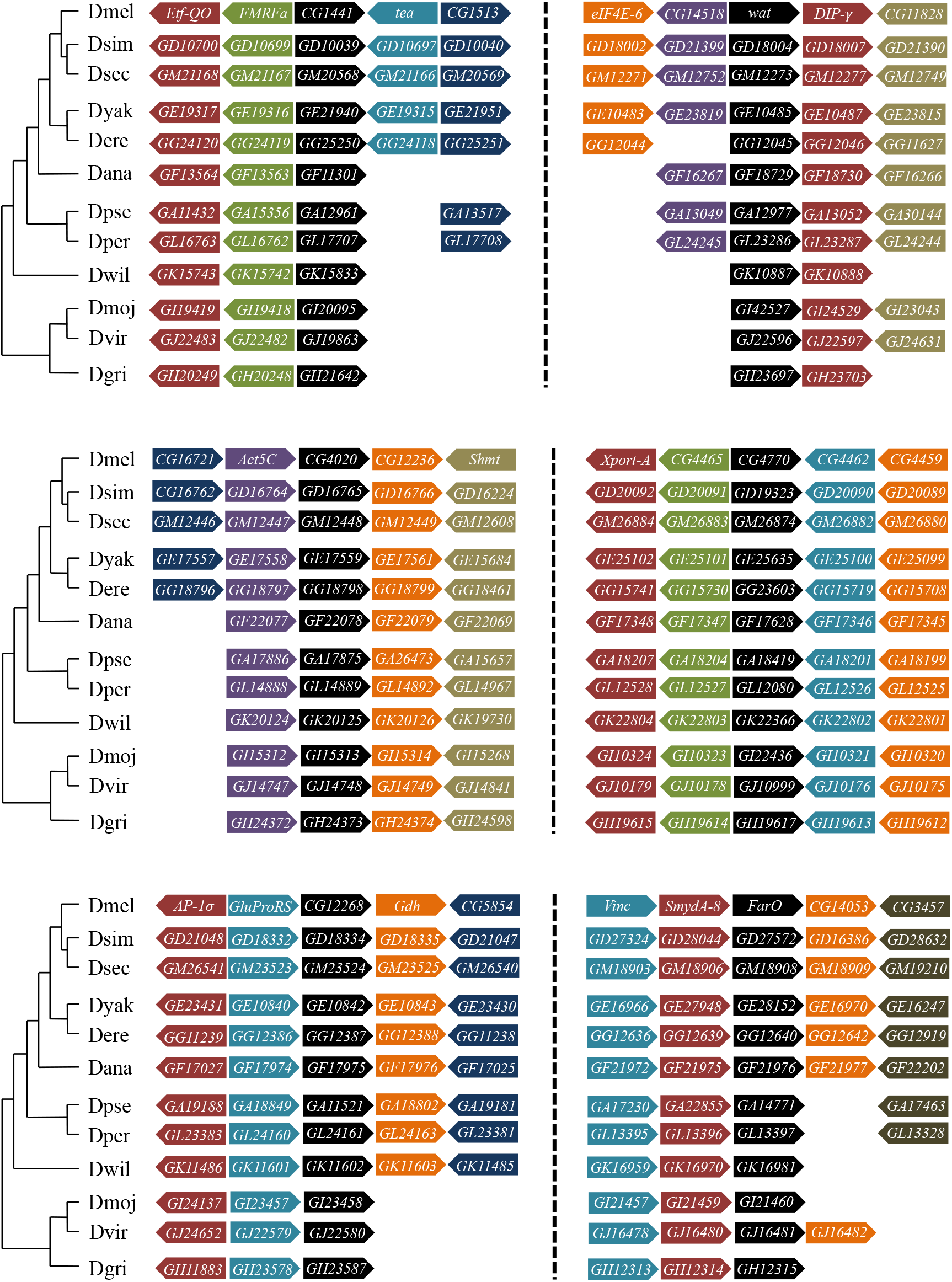
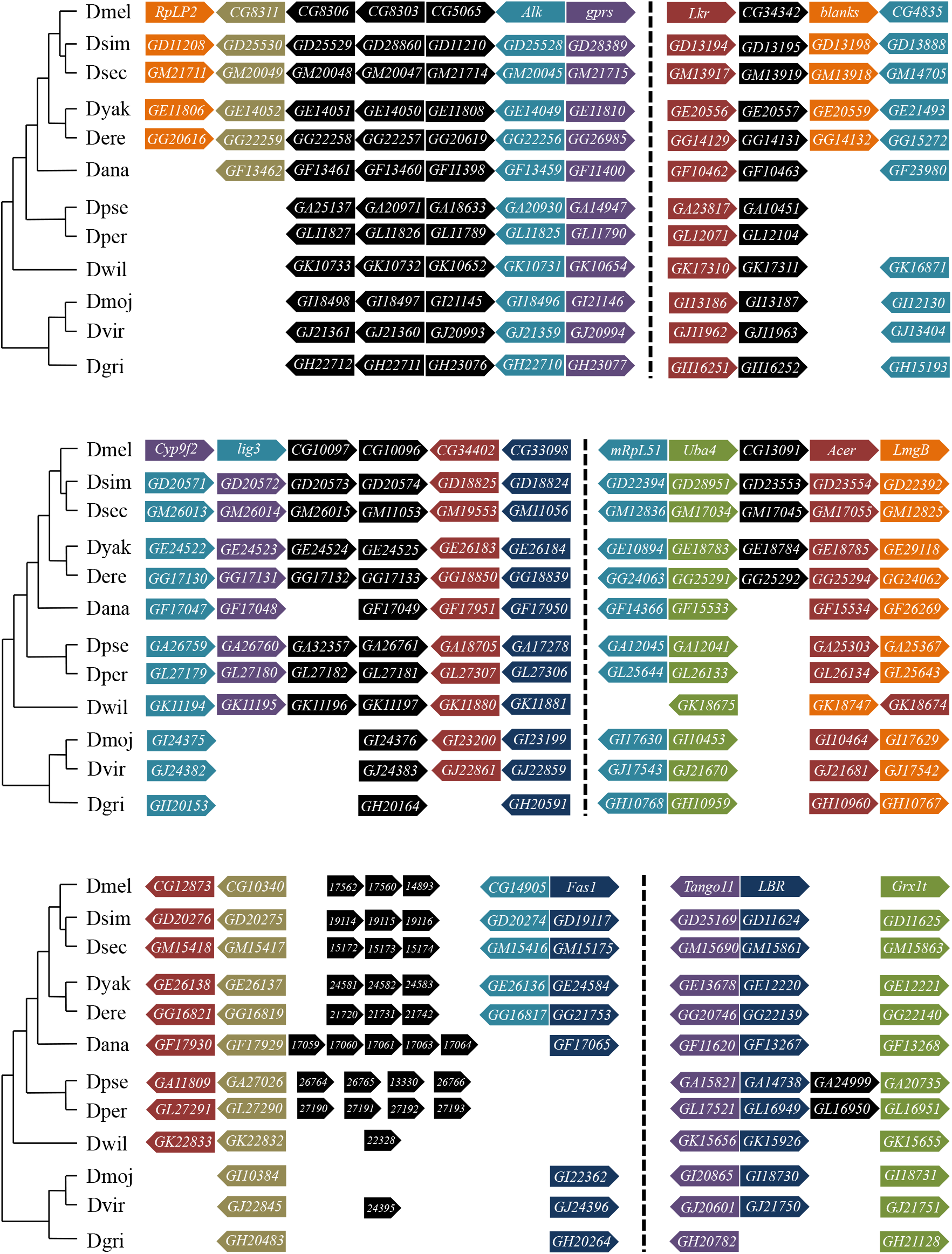
Synteny conservation at FAR genomic loci and adjacent genes in the *Drosophila* genus.

**Fig. S3.**
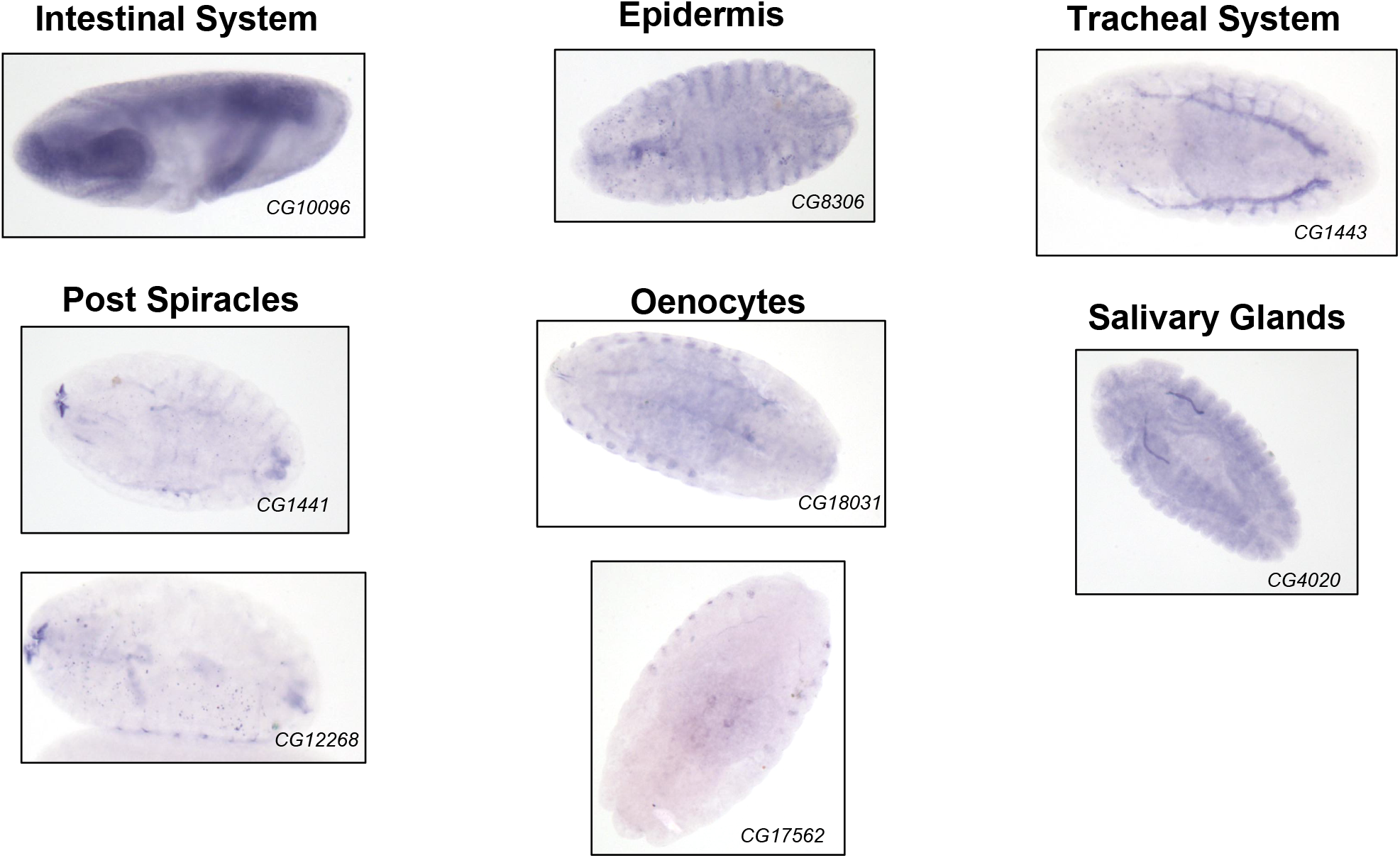
Expression of some of the FARs in *D. melanogaster* embryos.

**Table S1.**
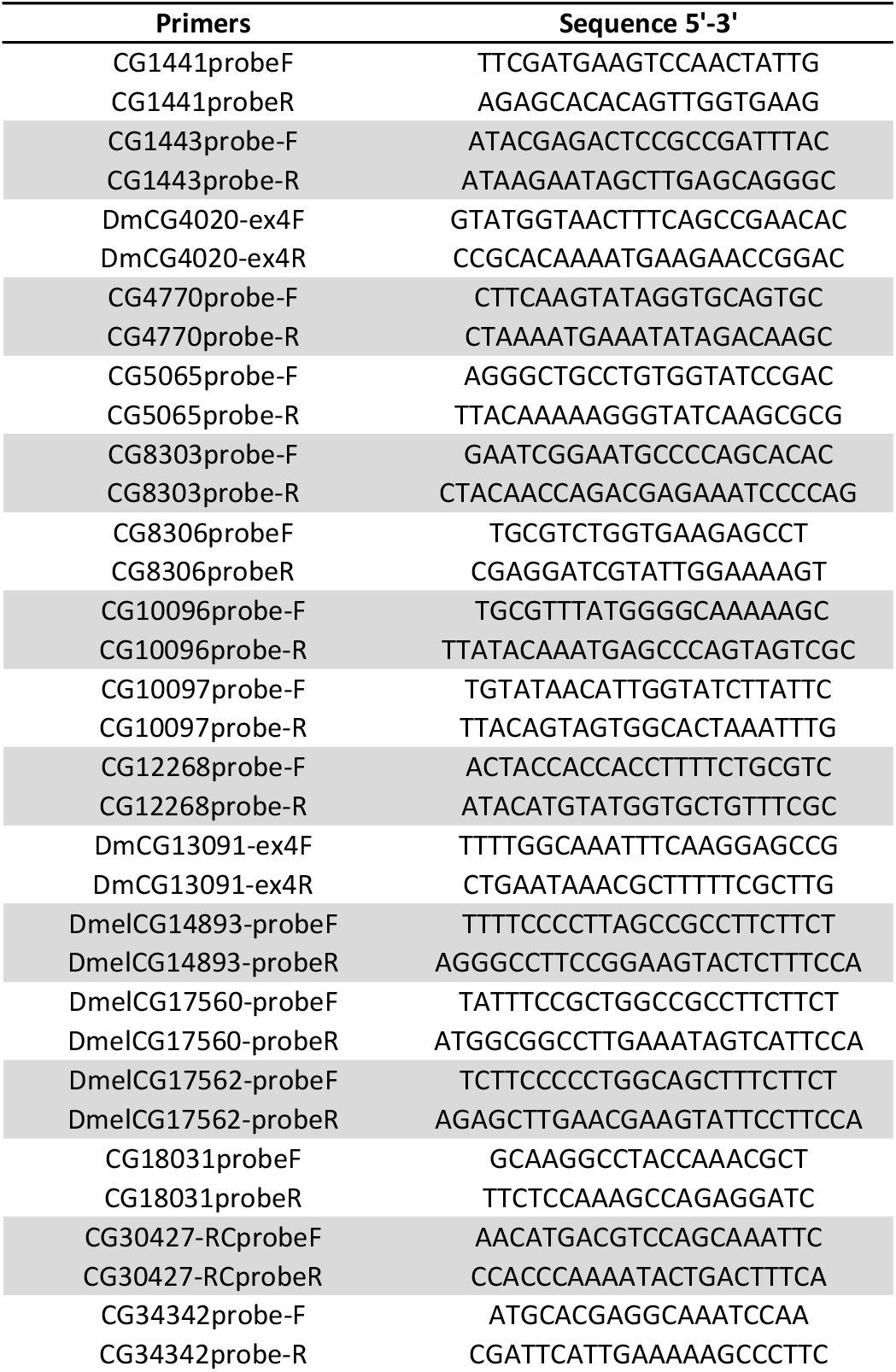
List and sequence of primers used in this study.

**Table S2.** Fly RNAi lines used in this study.

| Gene name | VDRC-ID | ON targets | Off targets |
| --- | --- | --- | --- |
| CG1441 | GD4034 | 1 | 1 (CG14162) |
| CG1443 | KK105518 | 1 | 0 |
| CG4020 | KK107095 | 1 | 0 |
| CG4770 | KK101641 | 1 | 0 |
| CG5065 | KK108205 | 1 | 0 |
| CG8303 | KK103744 | 1 | 0 |
| CG8306 | KK113348 | 1 | 0 |
| CG10096 | GD50765 | 2 (CG10097) | 1 |
| CG10097 | GD6089 | 2 (CG10096) | 1 (CG16974) |
| CG12268 | GD1166 | 1 | 0 |
| CG13091 | KK106304 | 1 | 0 |
| CG14893 | KK100324 | 1 | 0 |
| CG17560 | KK104756 | 1 | 2 (CG10096/CG10097) |
| CG17562 | GD37365 | 1 | 2 (CG10096/CG10097) |
| CG18031 | KK107564 | 1 | 1 (CG31019) |
| CG30427 | KK102330 | 1 | 0 |
| CG34342 | KK104554 | 1 | 2 (CG12109, CG10936) |

